# Longitudinal blood microsampling and proteome monitoring facilitate timely intervention in experimental type 1 diabetes

**DOI:** 10.1101/2024.11.19.624292

**Authors:** Anirudra Parajuli, Annika Bendes, Fabian Byvald, Virginia M. Stone, Emma E. Ringqvist, Marta Butrym, Emmanouil Angelis, Sophie Kipper, Stefan Bauer, Niclas Roxhed, Jochen M. Schwenk, Malin Flodström-Tullberg

## Abstract

Symptoms of immune-mediated diseases (IMIDs) typically appear after irreversible tissue damage, making early interventions based on pre-symptomatic indicators crucial. Current efforts to identify molecular markers of early disease lack the resolution, convenience and cost efficiency required to prevent irreversible tissue damage. Analyzing frequently self-collected samples, such as dried blood spots (DBS), could enable the earlier detection of diseases, identify disease-predictive markers and facilitate tailored interventions. To test this, we regularly microsampled a mouse model infected with a type 1-diabetes (T1D)-associated virus. This longitudinal DBS sample collection was analyzed for 92 circulating proteins, revealing transient molecular changes in virus-infected animals that would have been missed with less frequent sampling. Machine learning predicted infection status after day 2 post-infection with >90% accuracy, enabling well-timed treatment of virus-infected animals and diabetes prevention. Our study demonstrates the utility of frequent blood microsampling to monitor disease during the pre-symptomatic phase, allowing for timely interventions.

**Teaser:** Frequent blood microsampling detects early biomarkers, enabling timely intervention in immune-mediated diseases

## INTRODUCTION

An essential goal of precision medicine is to prevent diseases before they manifest (1). This relies on the identification of predictive biomarkers and the development of methods to detect them in asymptomatic individuals. However, many existing molecular tools and approaches are not yet optimized or tailored for the precise molecular monitoring of at-risk individuals.

A global analysis of plasma proteins can offer valuable insights into the different stages of a disease, disease progression, and treatment responses. As such, advanced proteomic techniques may be leveraged for early disease detection, ideally through the use of easily accessible body fluids (2). Recent breakthroughs have enabled precise and high-throughput measurements of blood proteins (2, 3). These cutting-edge techniques allow comprehensive proteome analysis and can identify subtle disease-related or even predictive protein changes (4). Despite new advances which facilitate the search for proteomic disease signatures (5), blood sampling still necessitates visits to healthcare centers. This inconvenient procedure often results in sampling at a single time point or timespans between samples that sometimes amount to months or even years. Lengthy sampling intervals limit the ability to capture rapid and transient proteome changes associated with disease risk and progression. Overcoming this limitation is crucial for timely disease detection and personalized disease interventions.

Promising remote sampling methods, which could permit frequent self-sampling, include dried blood spots (DBS). DBS samples can be shipped by regular mail to advanced laboratories. We recently demonstrated that unsupervised volumetric DBS sampling by finger-pricking at home allows for the precise measurement of multiple circulating antibodies associated with COVID-19 (6). Using state-of-the-art proteomic tools, we showed that 10 µl of dried blood was sufficient to successfully quantify hundreds of circulating proteins (7). Based on these observations, we hypothesized that frequent quantitative DBS analysis could be used to discover circulating proteins or early proteome signatures that are predictive of disease onset.

Immune-mediated diseases (IMIDs) occur when the immune system mistakenly attacks the body’s own tissues (8). Symptoms typically arise after significant and often irreversible damage has occurred. Most of these conditions, especially those with an autoimmune component, are chronic and significantly impact a person’s quality of life. Examples of IMIDs include multiple sclerosis (MS), rheumatoid arthritis (RA), and type 1 diabetes (T1D). T1D is characterized by the loss of functional pancreatic beta cells, which leads to reduced insulin production. The autoimmune origin of T1D is intimated by, in over 95% of those who develop the condition, the appearance of autoantibodies (AAbs) reactive to the islets of Langerhans approximately six months to a few years before disease onset. Additionally, autoreactive T cells are detected in the pancreas before clinical symptoms manifest (9). Environmental factors appear to play a crucial role in mediating the risk of developing T1D, but the exact triggers behind the break in immunological tolerance to the beta cells remain unknown (10). Several studies have indicated that an enterovirus infection may precede the first appearance of islet autoantibodies (11). Consequently, a well-timed antiviral therapy could intervene in the events that initiate islet autoimmunity and could also help to preserve beta cell function at disease onset (12). Treatment with an anti-CD3 antibody (Teplizumab), which targets T cells, has been shown to delay disease onset for up to a couple of years in AAb-positive individuals that are close to developing clinical symptoms (13-15). Identifying individuals who have been infected by a potentially diabetogenic virus but have not yet developed islet autoimmunity, or those who are islet autoantibody positive and near disease onset, remains a significant and currently unsolved challenge.

Building on our previous work, which demonstrated that home-based sampling effectively monitors the blood proteome (7), we hypothesize that quantitative DBS can reveal circulating proteins or proteome signatures predictive of T1D onset. In this study, we used experimental models of T1D (16-18) to investigate whether frequent blood microsampling can capture changes in the circulating proteome that occur early in disease processes, enabling timely, molecularly informed interventions.

## RESULTS

### Frequent blood microsampling and pancreas pathology in CVB3-infected NOD mice

To study the effects of infection with an enterovirus associated with T1D in humans (11) on the circulating proteome, two independent longitudinal studies (here denoted Study 1 and Study 2) were conducted in which female non-obese diabetic (NOD) mice, aged 8-9 weeks, were infected with Coxsackievirus B3 (CVB3) or mock-infected with a buffer. Previous studies have shown that the pancreas is permissive to CVBs, but infection of NOD mice in this age range does not induce diabetes (16, 17). For each mouse, 5 µl of blood was collected before infection and at several time points thereafter until 14 days post-infection (p.i.) (**Fig. 1A**). Each blood sample was collected on a filter paper disc (Capitainer). After the blood had dried, the DBS were stored at room temperature until analysis. Health measurements revealed that in both studies, the infected mice lost weight from days 5-6 p.i. (fig. S1). A few infected mice had to be euthanized on days 9 and 10 p.i. due to excessive weight loss (>15%, see fig. S1). Upon termination, the pancreases of buffer-treated animals appeared normal, while most infected mice had smaller pancreases with clear histopathological signs of infection (fig. S2). Notably, none of the animals developed diabetes (data not shown).

**Figure 1.**
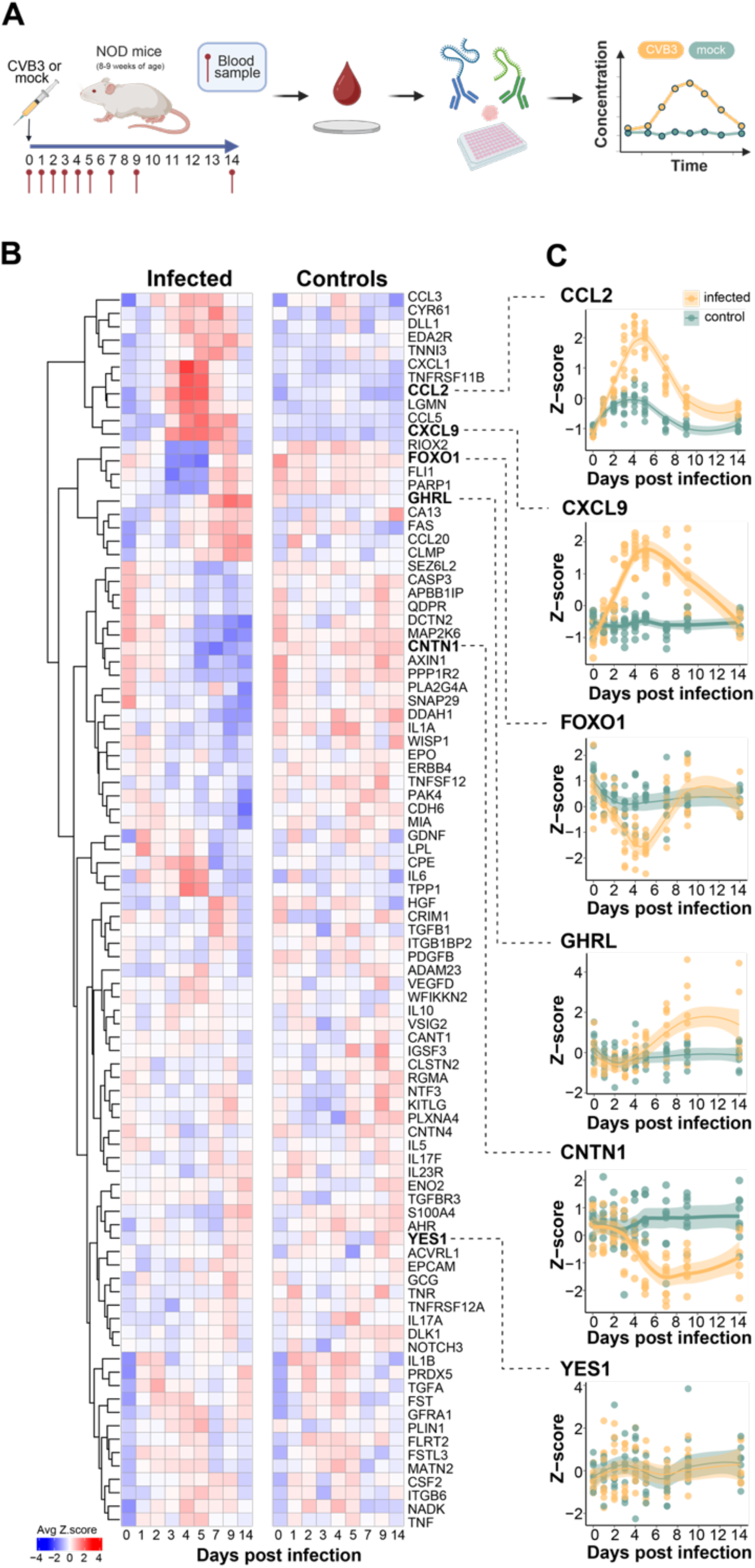
Longitudinal DBS protein profiling reveals dynamic proteome alterations in CVB3 infected NOD mice. **(A)** Schematic of study design. In two independent studies, NOD mice aged 8-9 weeks were infected with CVB3 (200 µl RPMI medium containing 10^5^ PFU CVB3, i.p., n = 9) or mock infected (200 µl of RPMI medium, n = 8). A blood sample (5 µl) was collected from the tail vein at indicated time points (see schematic) and dispensed onto filter discs (Capitainer). Blood samples were eluted and proteins were measured using proximity extension assays (Olink), followed by data analysis (**B**) Heatmaps showing the average protein levels (z-score) per sampling day for each protein for the infected (left) and control (right) groups. The proteins are clustered based on the levels in the infected group. Red indicates a high z-score and blue indicates a low z-score. (**C**) Protein profiles for six proteins. Each dot represents a sample, and color indicates if the included mouse belonged to the infected (yellow) or control (green) group. Smooth lines have been fitted to each group, with the 95% confidence interval around the smooth line.

### Proteome profiling of DBS samples collected from CVB3-infected NOD mice

DBS samples were analyzed using proximity extension assays (PEA). A total of 92 proteins were measured in longitudinal DBS samples from all animals. The data was first normalized using Protein-specific Probabilistic Quotient Normalization (ProtPQN), as previously described (7). Potential outliers were assessed, but none were identified (fig. S3A). The data was then scaled and centered using z-score transformation. After pre-processing, no significant differences in the overall protein levels were found between the two studies (p > 0.05), and the percent variance explained by the first principal component (PC1) was decreased from 80% to 17% (fig. S3B-D). Unless otherwise specified, the pre-processed and transformed data from the two studies were combined and used for the downstream analyses.

To assess the reproducibility of the affinity proteomics method, we concurrently collected multiple samples (n = 6 in Study 1, n = 5 in Study 2) from the same animal. Using pre-processed data, we observed an average coefficient of variation (CV) of 10.2% across all proteins in Study 1 and an average CV of 20.5% for Study 2. Per sample, Spearman correlations of rho = 0.99 for Study 1 and rho = 0.94 for Study 2 were obtained for the inter-animal replicate samples (fig. S4). This suggested that the data was suitable for further investigations.

The detectability of the proteomics data was calculated for each study using unprocessed NPX values by calculating the number of data points above the limit of detection (LOD). In study 1, 64.2% of data points were above the LOD, compared to 61.1% in Study 2. This detectability did not differ significantly between the infected and control groups (fig. S5A-B). When divided by sampling day, in Study 2 a greater number of proteins were detected above the LOD in the infected group on day 3 p.i., whereas the detectability was lower on days 1, 9, and 14 p.i. in this study (p < 0.05) (fig. S5D). No significant differences were found in Study 1 (fig. S5C).

### Longitudinal proteome profiling of DBS samples reveals dynamic and transient protein changes induced by CVB3 infection

To gain a first, unbiased overview of the temporal levels of circulating proteins, we performed hierarchical clustering of proteins over time and in response to the virus infection (**Fig. 1B,C**). The obtained protein clusters were then applied to the control data to illustrate the post-infection protein dynamics.

For example, in the infected group of animals, proteins involved in immune responses such as CCL2, CXCL9, CXCL1, TNFRSF11B, LGMN, and CCL5 showed an increase, peaking around day 4 p.i. Additionally, a noticeable yet minor increase in CCL2 levels was observed in the control group in the first 5-6 days p.i. (**Fig. 1B,C**).

Other proteins, including CCL20, GHRL, and CLMP, had an increased abundance in CVB3-infected mice from days 5-7 p.i. A cluster of proteins, including PARP1, RIOX2, FOXO1, and FLI1, decreased in level in infected mice between days 4-7 p.i. but returned to baseline by day 14. Conversely, protein levels for DCTN2, AXIN1, CNTN1, and CDH6 declined around day 5 p.i. and remained lower throughout the remaining sampling days.

The control group displayed a more uniform distribution of z-scores, with fewer changes in protein levels compared to the infected group of mice. Other proteins, like the protein YES1, remained unchanged over time in both groups of mice (**Fig. 1B,C**). An interactive web interface was developed to enable browsing of protein-centric results (URL will be provided upon publication). The interface presents the profile of each protein and summarizes the data in terms of precision, variance, and protein-protein correlations (fig. S6).

Next, we performed statistical tests to compare the protein levels in infected mice with those in the control group at each time point to identify infection-associated changes in protein levels (**Fig. 2**). Notably, CCL2 was significantly elevated in the infected group of animals compared to the uninfected control animals as early as 2 days p.i. (FDR adjusted p-value < 0.05), and this increase persisted until day 14 p.i. (**Fig. 2;** Data S1). Similarly, CXCL9 levels were elevated in the infection group from day 3 until day 9 p.i., indicating a temporal pattern in the host immune response. The highest number of proteins (n = 12) with significantly different time-dependent profiles between the infected and control animals was found on days 4 and 5 p.i. Seven proteins were elevated in the infected group on day 4 p.i., and six proteins were elevated in the same group on day 5 p.i. By day 14 p.i., significant differences were detected in the levels of six proteins when comparing the groups, with CCL2 being the only protein slightly elevated in the infected animals compared to the control group (**Fig. 2**).

**Figure 2.**
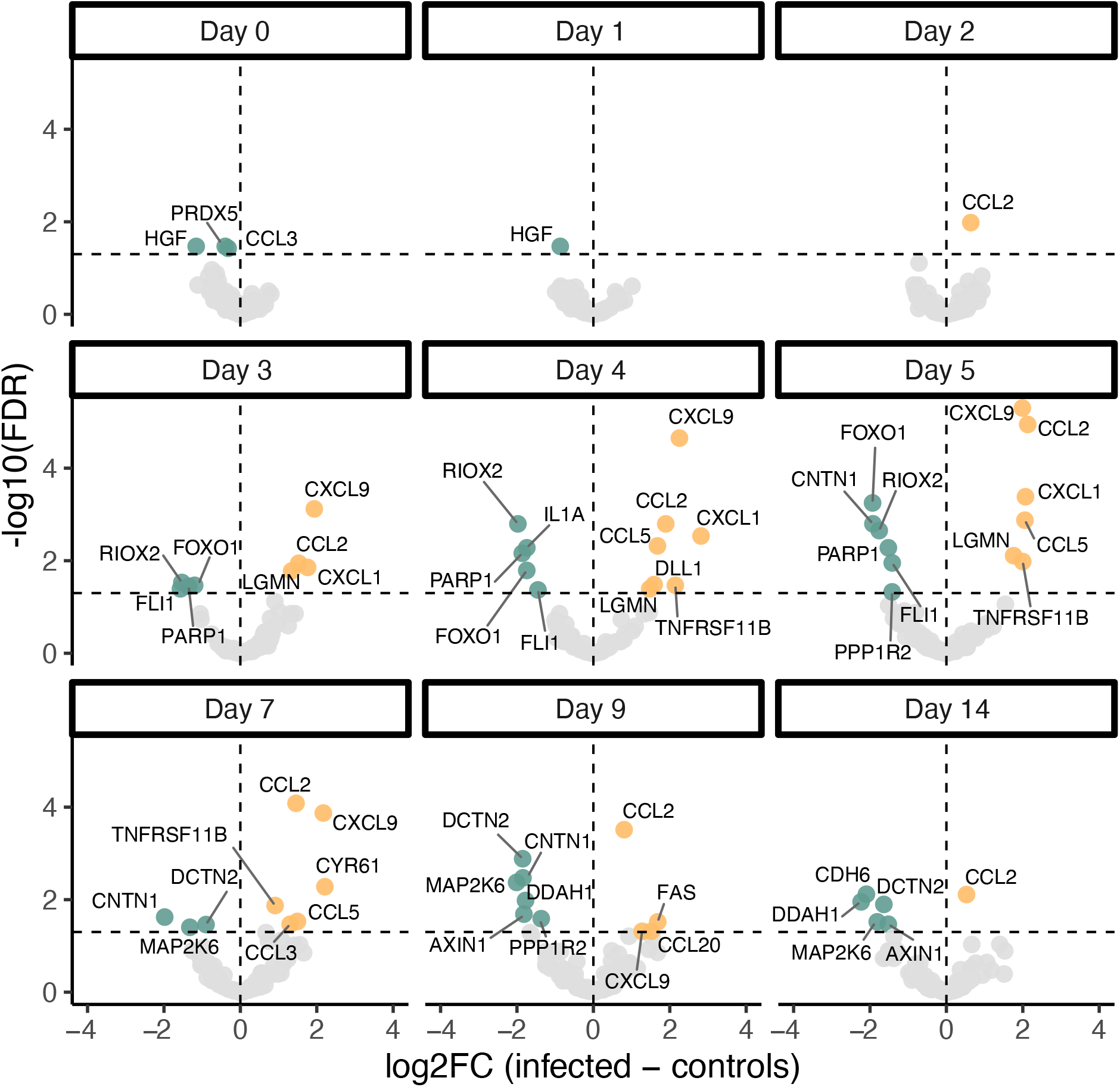
Protein dynamics across sampling time in CVB3 infected and control NOD mice. DBS samples were collected from CVB3- and mock-infected NOD mice for 14 days p.i. as described in Fig. 1A. Proteins were measured using proximity extension assays and the protein signals were transformed to z-scores before combining the data from two independent studies. The figure shows volcano plots for each sampling day for the log^2^-fold change between the infected and control groups plotted against the FDR-adjusted p-value obtained from two-sided Students t-test. Each dot represents a protein. The horizontal dotted line represents a p-value of 0.05, and proteins above are considered significant. The vertical dotted line represents a fold-change of zero, and proteins to the right of the line are found at higher levels in the infected group and vice versa. Yellow and green dots are proteins that are up- or downregulated respectively in the infected mice compared to the controls. Gray dots are proteins that were not considered significantly different between infected and control animals.

We also used generalized additive models (GAMs) to assess linear and non-linear relationships between protein levels and the number of days post-infection. This revealed a notable number of non-linear relationships between time post-infection and protein levels for several proteins after Bonferroni correction (n = 37 for the infected group, n = 8 for the control group; Table 1; Data S2). These included CCL2, CXCL9, and CXCL1, indicating that the level of these proteins fluctuated over time. Seven proteins (TNFRSF11B, CCL5, CCL2, CXCL9, CXCL1, GHRL, and EDA2R) had significantly different time-detection profiles between the groups using this model.

**Table 1.**
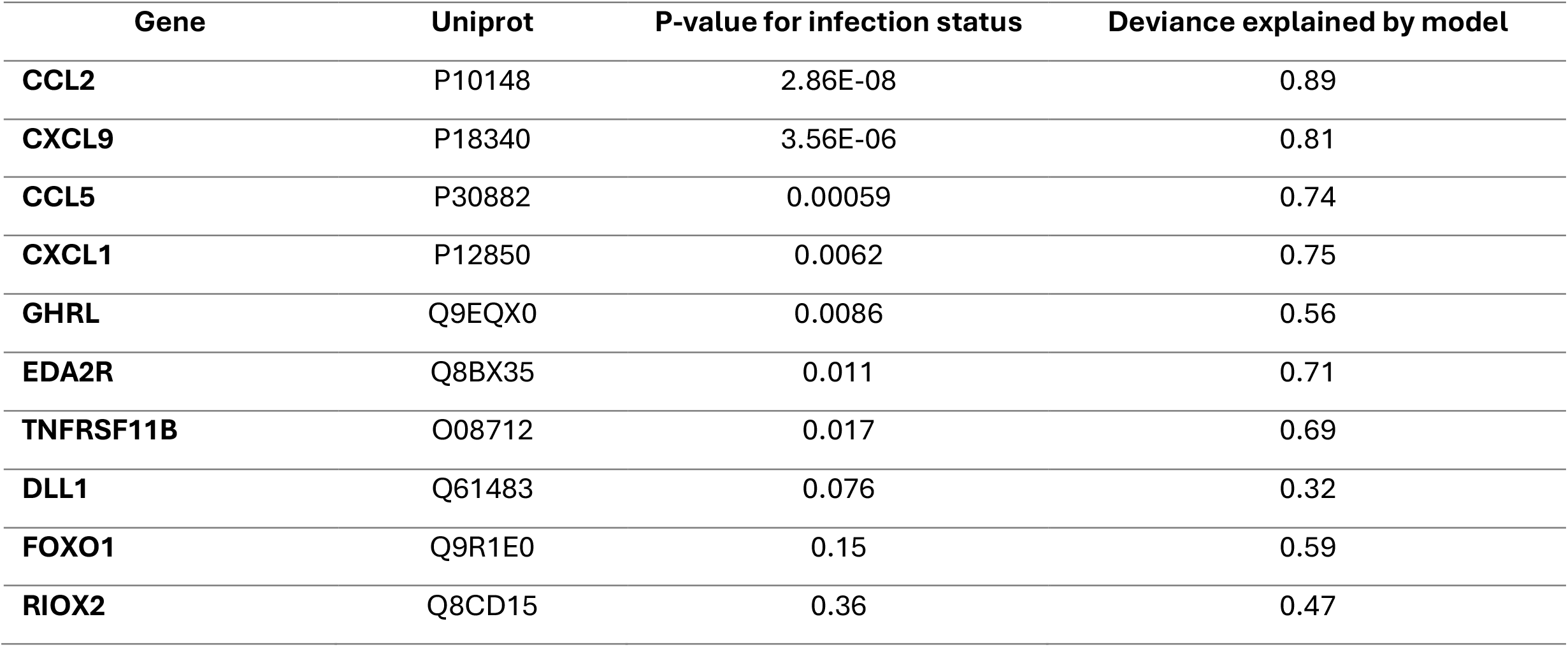
P-values from generalized additive model (GAM) for top 10 proteins differentiating CVB3 infected and control animals.

| Gene | Uniprot | P-value for infection status | Deviance explained by model |
| --- | --- | --- | --- |
| CCL2 | P10148 | 2.86E-08 | 0.89 |
| CXCL9 | P18340 | 3.56E-06 | 0.81 |
| CCL5 | P30882 | 0.00059 | 0.74 |
| CXCL1 | P12850 | 0.0062 | 0.75 |
| GHRL | Q9EQX0 | 0.0086 | 0.56 |
| EDA2R | Q8BX35 | 0.011 | 0.71 |
| TNFRSF11B | O08712 | 0.017 | 0.69 |
| DLL1 | Q61483 | 0.076 | 0.32 |
| FOXO1 | Q9R1E0 | 0.15 | 0.59 |
| RIOX2 | Q8CD15 | 0.36 | 0.47 |

### The levels of DBS proteins predict infection status early after infection

By applying machine learning (ML) techniques to our longitudinal proteome dataset (**Fig. 3A**), we next aimed to construct a classifier for infection prediction. Through preliminary analyses using the dataset generated in Study 1 and by applying permutation feature importance we identified the proteins CCL2 and CXCL9 as the most informative biomarkers for this task. Utilizing a multi-layer perceptron (MLP) classifier, we leveraged the trajectories of these proteins from day 0 to various time points p.i. (see Materials and Methods for detailed description). By day 2 p.i., we accurately predicted the infection status for 13 out of 17 mice (76%) and by day 3 p.i. 16 out of 17 (94%) (**Fig. 3B** and Data S3). For the one remaining mouse, which exhibited a delayed response trajectory, the infection status was correctly predicted by day 4 p.i. Overall, after day 2 p.i., our classifier correctly predicted the infection status for 83 out of 85 mouse-timestep combinations, equating to an accuracy rate of approximately 97.6%. A receiver operating characteristic (ROC) curve was calculated for the binary classification of infection status (**Fig. 3C)**, and the area under the ROC curve (AUROC) was 0.995.

**Figure 3.**
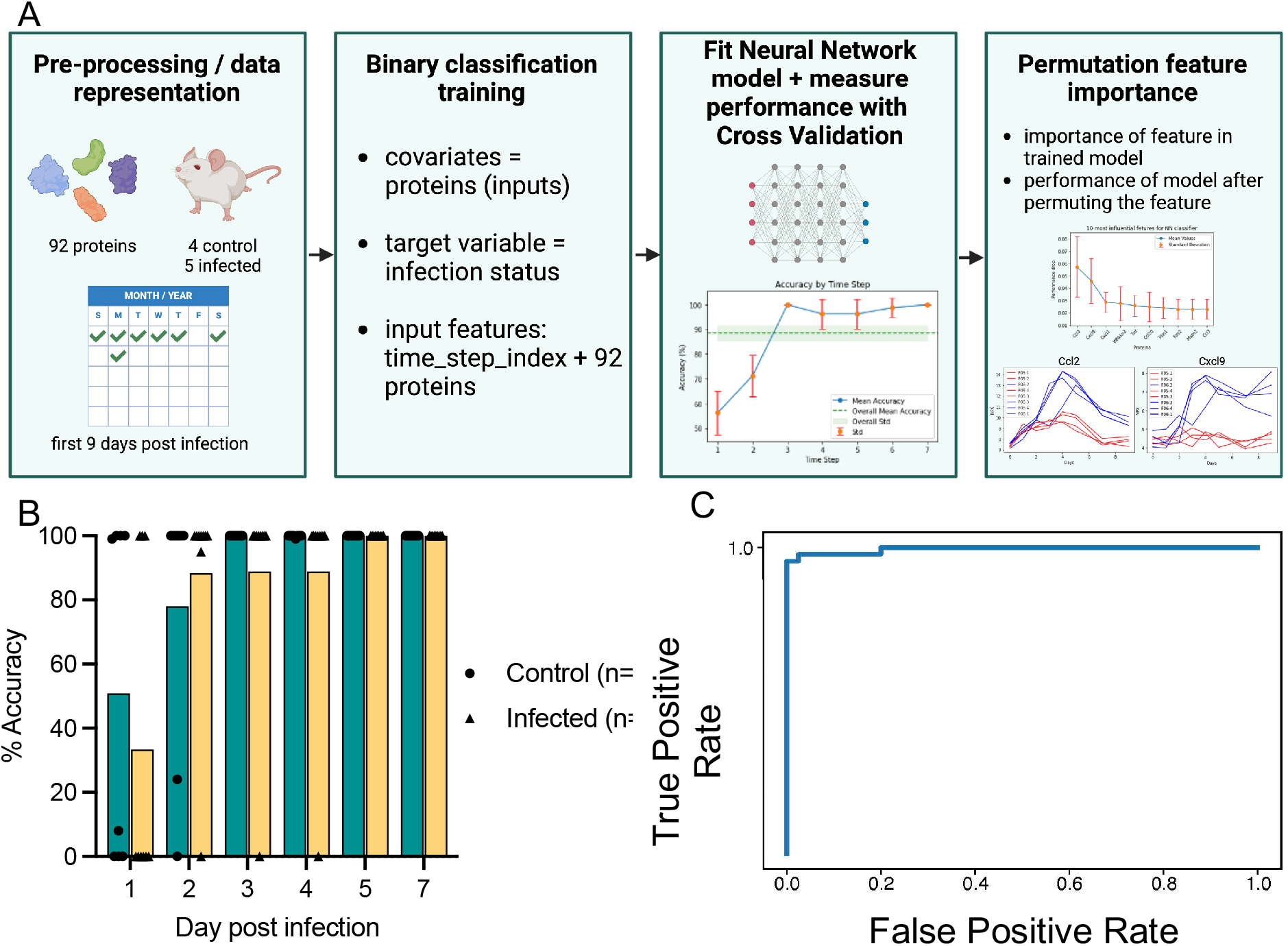
Model performance for classifier to predict infection status. (A) Machine learning pipeline for preliminary analysis of Study 1 samples aimed at identifying key features for infection prediction. Permutation feature importance was applied to quantify the contribution of covariates (proteins) to the model’s output. Here the model is a binary classifier that takes all 92 protein measurements at each time step as inputs for predicting infection status. Feature importance is assessed by randomly permuting each feature’s values and measuring the resulting drop in performance compared to the original unaltered input. Notably, permuting CCL2 and CXCL9 resulted in the largest accuracy drops, underscoring their significance. Their predictive relevance is visually confirmed in the plot depicting their temporal evolution. Note that the model used in the preliminary analysis differs from the one employed in the main experiments. In the “Fit Neural Network” box, step 1 = day 1, step 2 = day 2, step 3 = 3, step 4 = day 4, step 5 = day 5, step 6 = day 7 and step 7 = day 9. (B-D) Results from the model used in the main analysis including data from both independent studies (Study 1 and Study 2). (B) Percentage accuracy of the model up to day 7 post infection in control (n=8; green; circles) and infected (n=9; yellow; triangles) mice. Individual animals are represented by individual symbols. (C) Receiver operating characteristic (ROC) curve illustrating the performance of the binary classification for infection status. Sensitivity (true positive rate) is plotted against 1-specificity (false positive rate). The AUROC was 0.995.

### Monitoring-informed intervention prevents virus-induced diabetes development

The ML-based classifier demonstrated that infection status could be accurately predicted at an early point p.i. (day 2), even before clear signs of infection, such as weight loss (fig. S1) became noticeable. Therefore, we next explored whether this early disease prediction could offer a window for preventive treatments.

To this end, we employed an experimental model for CVB-induced T1D, the SOCS-1-tg mouse model (16-18). In these NOD mice, the overexpression of the suppressor of cytokine signaling-1 (SOCS-1) under the control of the insulin promotor renders the beta cells unresponsive to interferon (IFN) stimulation. Consequently, critical autonomous antiviral defense mechanisms, typically activated during virus infections via type I IFNs, are compromised in the beta cells. As a result, the beta cells succumb to CVB infection, and SOCS-1-tg mice develop diabetes within approximately 5-12 days of infection (16-18).

While there is a lack of potent antiviral treatments for CVB infections, passive immunizations have shown promise (19-21). To explore this approach, we generated non-immune and immune sera from a separate group of NOD mice. Subsequently, SOCS-1-tg mice were infected with CVB3, followed by treatment with non-immune or immune sera on days 2 and 3 p.i. (as depicted in **Fig. 4A**). In the non-immune sera-treated group, diabetes was triggered in 5 out of 8 mice (**Fig. 4B-C**). A histological assessment of pancreas specimens revealed that most of the animals suffered from extensive exocrine tissue damage and a minimal number or absence of insulin-positive cells in the islets (**Fig. 4D-E**, fig. S7). Remarkably, among the 6 mice treated with immune sera during the same time frame, none (0/6) developed diabetes (**Fig. 4B-C**) and their pancreases remained intact with islets expressing insulin and glucagon (**Fig. 4D-E**, fig. S7).

**Figure 4.**
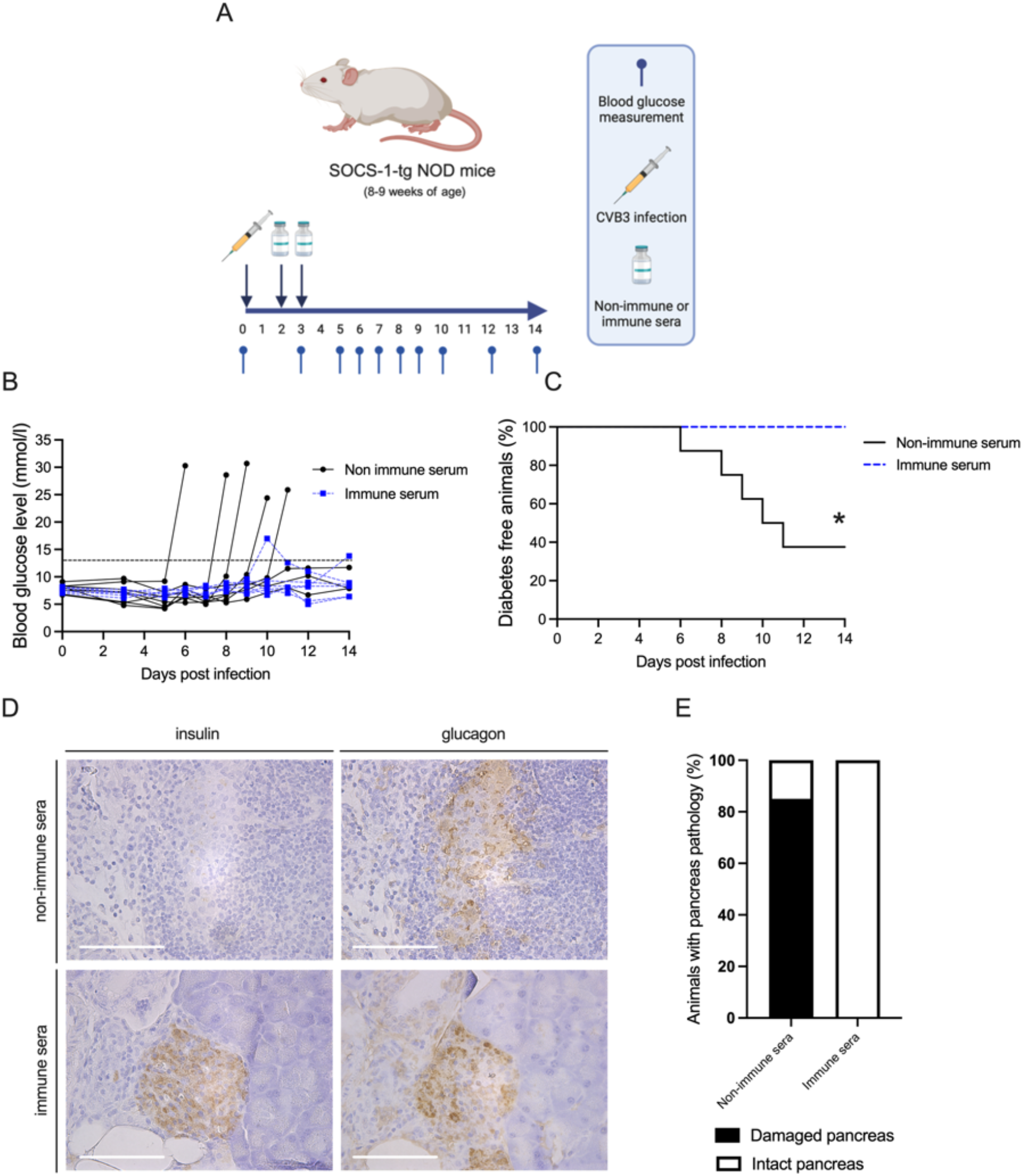
Early intervention prevents virus-induced T1D. SOCS-1-tg mice were infected with CVB3 (10^5^ PFU CVB3, i.p.). On day 2 and 3 p.i., animals were treated with either non-immune (n = 8) or immune sera (n = 6) by i.p. injection (total volume 200 µl/mouse) as shown in the experimental schematics in (**A**). (**B**) Blood glucose values of individual animals treated with non-immune sera (black line, n=8) or immune sera (dotted blue line, n=6). Mice were deemed diabetic when the blood glucose level was equal to or exceeded 18 mmol/l, or when two consecutive daily measurements ranged between 13 and 18 mmol/l. Dotted black line marks 13 mM glucose. (**C**) Diabetes incidence curves summarizing results shown in (B). *p<0.05 comparing the two groups by Log-rank (Mantel-Cox) test. (**D**) Representative images of sequential pancreas sections from mice infected with CVB3 and treated with non-immune or immune sera. Pancreas was stained for insulin (panels on left) or glucagon (panels on right). Positive areas are stained brown. Scale bars = 100 µm. (**E**) Percentage of animals with damaged and intact tissue morphology in pancreas specimens from mice treated with non-immune sera (n = 7) or immune sera (n = 5). The infection status of two animals, one from each treatment group, could not be determined histologically due to insufficient quality of the FFPE sections, and were excluded from the assessment.

## DISCUSSION

To achieve precision medicine for the preventive treatment of IMIDs, identifying early, non-symptomatic disease stages is crucial. While sensors and apps can help track symptoms, conditions and/or behavior, there is still a lack of tools to monitor molecular changes that can serve as early indicators of disease. Of the accessible sample types, blood provides system-wide and organ-specific insights into health. Proteins, lipids, and other small molecules can be reliably measured in large numbers to track changes (22). However, the timing of blood draws is crucial for identifying changes before the tipping points are reached, making longitudinal sampling in pre-symptomatic individuals necessary.

In recent years, thanks to the development of blood self-sampling using DBS, new routines for health monitoring have been introduced (23). The ease of fingertip blood collection and the advent of technologies compatible with small sample volumes raises the question of whether the data collected after frequent sampling can effectively track disease progression and enable early preventative interventions. Using mouse models and focusing on proteins identified in blood samples, we asked if frequent blood microsampling and the analysis of time-resolved protein patterns could identify predictive disease markers and the optimal window for effective disease interventions. We demonstrate that repeated sampling of small amounts of blood (5 µl), followed by multiplexed protein analysis of DBS samples, can efficiently monitor proteome changes during an infection with a virus linked to T1D and enable timely disease intervention.

By analyzing the DBS samples using a commercially available protein panel, we monitored > 90 proteins relevant to various biological processes and disease areas. These included immune response proteins, growth factors, cell signaling proteins, proteins involved in cardiovascular and neurological diseases and proteins associated with various cancers. Among the immune response proteins, the chemokine CCL2 showed a transient increase in infected and control animals. In CVB3-infected animals, CCL2 levels peaked around day 5 p.i., while the peak was earlier and much lower in the control group. This indicates that the initial blood sampling triggered CCL2 production in both groups, as CCL2 plays a crucial role in the recruitment of monocytes and other immune cells to sites of inflammation caused by tissue injury or infection (24). Minor but similar temporary increases in other immune-related proteins, such as CCL20 and IL1β, were detected in both groups (**Fig. 1**). This observation is important to consider, as it demonstrates that the sampling procedure itself, especially in circumstances where samples are repeatedly taken from the same location, may temporarily impact the levels of certain proteins. The protein data from the uninfected control group, however, displayed a more uniform distribution of z-scores with fewer protein level changes than the infected group (**Fig. 1**). This suggests that, overall, the mice tolerated the repeated handling and blood sampling well.

Recent work has summarized the current understanding of the immune response to CVB infection (11). Our study offers a high resolution of systemic proteome alterations during the acute phase of the disease. We observed the highest number of altered circulating proteins between days 4 and 7 p.i. Despite being underrepresented in the panel, most of these proteins were associated with the immune system. Given that robust innate antiviral defense mechanisms are activated and peak earlier than day 4 p.i. (20, 25), future studies using a wider panel that covers additional immune-related proteins may identify further candidates with altered abundance early p.i. Complementary insights into infection biology could also be provided by other proteomics methods, such as mass spectrometry, as well as the analysis of other molecules including metabolites and/or lipids. Our study underscores the dynamic nature of protein changes in response to acute infection, with specific proteins exhibiting significant temporal patterns. Importantly, these patterns would have been missed with less frequent sampling.

The dynamic protein changes observed during acute CVB infection also revealed proteins that decreased in abundance. For example, FOXO1 and CNTN1 levels temporarily dropped during infection. The implications of these changes are an area of future interest. Interestingly, there was a trend towards a late increase in ghrelin levels in CVB3-infected mice. Ghrelin, known for stimulating appetite and regulating hunger and energy balance, typically increases after significant weight loss (26). The CVB3-infected animals experienced weight loss throughout the infection, likely due to extensive damage to their exocrine pancreas tissue. Collectively, this highlights that through the analysis of longitudinal DBS samples, we can identify the molecular footprints of the direct biological consequences of CVB3 infections, as well as their indirect, broader impacts on health.

ML methods are becoming pivotal for analyzing complex datasets, particularly for the prediction and staging of diseases, and for individualized patient care (2). After applying ML techniques to our longitudinal proteome dataset, we successfully constructed a classifier for infection prediction. The proteins CCL2 and CXCL9 emerged as the most informative biomarkers for this task, consistent with classical approaches. By day 2 p.i. we accurately predicted the infection status with an accuracy rate of > 75% and by day 3 p.i. at a rate of > 95%. The effectiveness of this prediction for enabling timely interventions was demonstrated using an experimental model for virus-induced T1D (16-18). Infected animals treated with immune sera on days 2 and 3 p.i. were protected from significant pancreatic damage and the development of T1D. These results underscore the potential of ML-based approaches in utilizing plasma proteomics for early and precise infection status prediction, opening new avenues for appropriate interventions and improved disease management.

Prospective cohort studies which have followed children from birth have been highly informative in enhancing our understanding of the pathogenesis of T1D (27-30). However, these studies have not yet identified the early triggers of beta cell autoimmunity and diabetes development. This limitation may be partly due to the relatively lengthy sampling intervals (e.g. three months or more between samples). Our proof-of-concept study highlights that more frequent sampling can help capture transient changes induced by a trigger. It also provides a more accurate picture of biomarker fluctuations, which may be useful for timely disease interventions using, for example, an antiviral or anti-CD3 treatment.

In conclusion, our proof-of-concept study underscores the importance of longitudinal sampling and molecular monitoring in elucidating the temporal dynamics of protein changes during the early stages of disease. This approach, validated in a mouse model, holds significant potential for identifying novel disease-predictive biomarkers and informing early intervention strategies for IMIDs such as T1D, MS, and RA.

## MATERIALS AND METHODS

### Animal husbandry and monitoring of animal health

NOD and SOCS-1-tg NOD (16, 31) mice were bred and maintained under specific pathogen free conditions at the Karolinska University Hospital Huddinge, Stockholm, Sweden. All experiments received approval from a local ethics committee (Linköpings djurförsöksetiska nämnd, Dnr 9222-2019 and 04291-2024) and were conducted in line with the NIH principles of Laboratory Animal Care, institutional guidelines at Karolinska Institutet and Swedish national laws. The animals were kept in ventilated cages with unrestricted access to food and water. Each cage housed a maximum of 5 mice, with no individual housing of animals. The animals were randomly allocated to different treatment groups. Mice underwent comprehensive health checks and changes in health status were monitored (including weight fluctuations, changes in natural behavior, porphyria, movement and posture, piloerection, respiration, and skin condition). The researchers were aware of the experimental groups throughout the experiments. At the end of the experiment, at a terminal time point deemed by weight loss (>15% of the maximum body weight during the experimental duration) or at the onset of diabetes, the mice were anesthetized with isoflurane, and then the mice were euthanized by cervical dislocation.

### Virus infections and serum transfers

Animals received an intraperitoneal injection of 10^5^ plaque forming units (PFU) of CVB3 (Nancy) or vehicle (RPMI media alone), in a total volume of 200 ml. Blood samples were collected at the indicated time points until day 14 p.i. To produce immune sera, male and female NOD mice aged 11-12 weeks were infected with CVB3 and blood was collected via terminal heart puncture performed on anesthetized animals two weeks p.i. Non-immune sera were produced from blood collected from age-matched uninfected NOD mice. Pooled and heat-inactivated sera (56°C for 30 min) were given to SOCS-1-tg NOD mice on day 2 and 3 p.i. (200 µl/mouse injected i.p.).

### Blood collection, glucose measuring, and monitoring of diabetes development

Blood was collected for proteome analysis from the outer tip of the tail. Prior to each collection the tail was wiped with 70% ethanol. On the first collection time point, a small portion (1 mm) of the tail extremity was removed using sterile scissors. A small droplet of blood (less than 10 µl) was extracted by gently holding the mouse by its tail. From this, 5 µl was pipetted onto a paper-based sampling disc (Capitainer AB, 710-0020) and left to dry until analysis. The scab was gently removed for subsequent blood collections. Blood glucose levels were determined using a Bayer Contour XT blood glucose meter (Bayer, Basel, Switzerland) on blood samples drawn from the tail vein. Diabetes was diagnosed when blood glucose levels were equal to or exceeded 18 mmol/l or when two consecutive daily measurements ranged between 13 and 18 mmol/l.

### Histology and immunohistochemistry

Pancreas specimens were fixed in formalin and embedded in paraffin. Five µm thick sections were cut using a Microm HM355S microtome (Thermo Scientific, Kalamazoo, MI, USA). A standard hematoxylin-eosin (H&E) staining was performed for the assessment of pancreas tissue morphology. Immunohistochemistry staining for insulin and glucagon was carried out as previously described (16, 32). The primary antibodies used were guinea pig anti-insulin (N1542, DakoCytomation. Glostrup, Denmark; 1:20,000 dilution) and rabbit anti-glucagon (ab92517, Abcam. Cambridge, UK; 1:24,000 dilution) and the secondary antibodies were biotinylated anti-guinea pig (DakoCytomation; 1:200 dilution) and biotinylated polyclonal anti-rabbit (Dako, Glostrup, Denmark; 1:500 dilution) for insulin and glucagon staining, respectively. Insulin and glucagon positivity were revealed by DAB staining, and sections were counterstained using hematoxylin. Stained tissue was examined, and images were acquired using a Leica DFC495 light microscope (Leica Microsystems Ltd. Heerbrugg, Germany) and LAS X software (v. 3.0.14.23224; Leica Microsystems Ltd.).

### Blood sample processing and proteomics assay

Filter discs with dried blood were randomized in flat-bottom 96-well plates (VWR, #10861-562) and eluted in 50 µl PBS with 0.05% Tween20 and protease inhibitor (cOmplete Tablets EASYpack, Roche, Ref#04693116001). After 1 h incubation with gentle shaking (170 rpm) at 23°C, the plates were spun at 400 g (Allegra X-12R, Beckman Coulter Inc) for 1 minute to remove any liquid from the plate seal. The supernatant was then transferred to a 96-well PCR plate and centrifuged at 2100 g (Allegra X-12R, Beckman Coulter Inc.) for 3 min. After centrifugation, 40 µl of the supernatant was transferred to a new 96-well PCR plate before starting the proteomics assay.

The Olink Target 96 Mouse Exploratory panel (Art # 95380; Batch B24933 for Study 1, Batch B30620 for Study 2) was used to measure 92 circulating proteins in the dried blood spot (DBS) eluates according to the manufacturer’s protocol for plasma samples. Briefly, 1 µl of each sample was incubated overnight with oligonucleotide-coupled antibody pairs, which only allows the hybridization of the DNA strands if the antibodies bind to a common protein in close proximity. In a subsequent step, proximity extension was used to create unique DNA reporter sequences for each protein and control, which were amplified and detected by real-time PCR using a Biomark™ HD (Fluidigm). The two sample batches were processed separately.

The Ct values were transformed into Normalized Protein eXpression (NPX) values using Olink NPX Signature (v. 1.6.0.0 for Study 1, v1.9.0 for Study 2). NPX is Olink’s arbitrary unit of relative protein levels and is reported on the log2 scale. The data sets of Study 1 and Study 2 were independently normalized per protein using ProtPQN to reduce sample-to-sample variation in DBS (33, 34). The presence of sample outliers was investigated by plotting the sample median against the interquartile range (IQR). A standard deviation (SD) of 5 was used as a cutoff for a sample being flagged as an outlier. The data was then scaled and centered by transforming it to z-scores per protein and per study.

### Statistical analysis

Statistical analyses were performed using Prism 10 software (version 10.3.1; GraphPad, La Jolla, CA) and R (version 4.4.1) unless otherwise stated. Plots were produced in Prism X or R using the ggplot2 package (version 3.5.1). Data are expressed as mean ± SD. False Discovery Rate (FDR) correction of multiple testing was performed using the Benjamini-Hochberg/Bonferroni method. A nominal p*-*value or FDR < 0.05 was considered statistically significant.

The reproducibility of the DBS sampling and the proteomics assay process was tested in biological replicates taken at the same timepoint (six samples from one mouse in Study 1, and five samples from one mouse in Study 2), and in technical duplicates. Spearman correlations between replicates were determined using the corr.test function from *psych* package (version 2.4.6.26) and visualized with *ggpairs* functions from the *GGally* package (version 2.2.1). The compatibility of the transformed data from the two studies was assessed using Wilcoxon rank sum test using the *wilcox*.*test* function from the *stats* R-package, and visualization using principal component analysis (PCA) plots. Heatmaps for the average protein expression for the infected and control groups per sampling day were generated using the *ComplexHeatmap* package in R (version 2.20.0). Proteins were clustered using the Euclidean distance matrix. Differences in detectability between infected and uninfected groups were investigated using Fisher’s exact test using the *fisher*.*test* function from the stats package. Two sample Student’s T-tests were used to determine differences in protein levels between infected and control mice at each time point.

Generalized Additive Models (GAMs) were used to assess the relationship between protein expression levels and infection status over time. The analysis used the *mgcv* package in R (version 1.9-1). A separate GAM was fitted for each protein assay, where the z-score was considered a function of infection status and sampling day with a smoothing term for sampling day with separate smooths for each infection status (Table 1, Data S2). The smoothness parameter (k) was set to 6 to control for potential overfitting. Repeated measures within subjects were accounted for by adjusting for subject-dependent variations in intercepts. The models were fitted using the Restricted Maximum Likelihood (REML) method to optimize the smoothing parameter. For p-values below 2e-16, the values were substituted with 2e-16 due to limitations in machine epsilon accuracy, as indicated in Table S1 and Data S2.

### Generation of a classifier to predict infection status

ML techniques were employed in python (version 3.7) to develop a binary classifier aimed at predicting the infection status of animals followed in the longitudinal study. Specifically, the classifier utilized the protein trajectories of each mouse from Day 0 until a given time step T to make its predictions. The two most informative proteins for this task, identified through preliminary analyses, were CCL2 and CXCL9, and their measurements served as the primary inputs for the model.

Each protein’s trajectory from Day 0 to Day T was encoded into a 4-tuple consisting of the following features:

Timestep T

- Standard deviation of protein measurements from Day 0 until Day T
- Range length of protein measurements from Day 0 until Day T
- Total variation of protein measurements from Day 0 until Day T

Where:

- Range length was defined as the difference between the maximum and minimum values of protein measurements from Day 0 to Day T.
- Total variation was the sum of the absolute values of the differences between consecutive measurements from Day 0 to Day T.

Thus, for each mouse, M, at timestep T, the representation is encapsulated in a 7-tuple: (T,std_devCcl2,range_lengthCcl2,total_variationC cl2,std_devCxcl9,range_lengthCxcl9,total_variati onCxcl9)

We chose a Multi-Layer Perceptron (MLP) for our classifier due to its simplicity and effectiveness in binary classification tasks (35, 36). The dataset comprised of 17 mice, each measured at 7 distinct time points in addition to Day 0, resulting in a total of 119 data points (17 mice × 7 time steps). A leave-one-out cross-validation approach was employed to validate the model. In this procedure, we iteratively selected one mouse for testing while training the classifier on the remaining 16 mice. This process was repeated 17 times, ensuring that each mouse was used as a test case once. To further ensure the robustness of our results, this cross-validation process was repeated 100 times.

A ROC curve for the binary classification of infection status was generated using the 17×5 output scores taken from the 17 classifiers trained with the leave-one-out cross-validation approach. Each classifier provided five predictions for the days following day 2 (i.e., days 3, 4, 5, 7, and 9).

### Web-based interface

The *shiny* package (version 1.8.1.1) was used to create an interactive web interface for the study. The app allows a protein-centric visualization of the proteomics data with interactive heatmaps for protein-protein correlations using *heatmaply* package (version 1.5.0). All packages and versions used for visualization are stated in the app.

## ACKNOWLEDGMENTS

We extend our gratitude to Ms. Selina Parvin from Karolinska Institutet, Stockholm, Sweden, for her invaluable assistance with histological analyses. We also thank the animal staff at the Preclinical Laboratory (PKL) Facility, Karolinska University Hospital Huddinge and Karolinska Institutet, for their support in breeding and housing the experimental animals. We thank Leo Dahl for all the fruitful discussions and the team at SciLifeLab’s Affinity Proteomics Unit in Stockholm for technical support.

Figures **1A**, **3A** and **4A** were created using Biorender.com.

Figure **1A**: Created in BioRender. Byvald, F. (2024) BioRender.com/t83q649

Figure **3A**: Created in BioRender. Byvald, F. (2024) https://BioRender.com/y07z021

Figure **4A**: Created in BioRender. Byvald, F. (2024) BioRender.com/b43t411

## FUNDING

This work was supported by grants from the: Swedish Child Diabetes Foundation (MFT) The Swedish Diabetes Foundation (MFT)

Karolinska Institutet, Sweden, including the Strategic Research Programme in Diabetes (MFT)

Swedish Research Council, grant numbers 2020-02969 (MFT)

Novo Nordic Foundation NNF18OC0034158 (MFT) Novo Nordic Foundation NNF24OC0092507 (MFT)

KTH Royal Institute of Technology, Digital Futures seed funding grant (SB, NR, JMS)

SciLifeLab’s Pandemic Laboratory Preparedness program VC-2021-0033 (JMS)

SciLifeLab’s Pandemic Laboratory Preparedness program VC-2022-0028 (JMS)

Swedish Research Council 2022-01374 (NR)

## AUTHOR CONTRIBUTIONS

Conceptualization: MFT, JMS, NR, SB Methodology: VMS, AB, MFT, NR, JMS, SB Investigation: AP, AB, FB, VMS, EA, EER, MB, SK Visualization: AP, AB, FB, EA, MFT, JMS Supervision: MFT, JMS, NR, SB Writing—original draft: MFT, AB, FB, JMS Writing—review & editing: AP, VMS, EER, MB, SK

## DATA AND MATERIALS AVAILABILITY

Upon publication, all data needed to evaluate the conclusions in the paper are present in the paper, the supplementary materials, and/or are accessible at URL (will be provided upon publication). Additional visualizations are presented in the Shiny app interface URL (will be provided upon publication). Codes used in this work will be made available upon publication via GitHub.

## SUPPLEMENTARY MATERIALS

Fig. S1. Percentage weight change in mock and CVB3-infected NOD mice.

Fig. S2. NOD mouse pancreas displays clear signs of pathology after CVB3 infection.

Fig. S3. Initial data analysis of proteomics data generated from dried blood spot samples collected from CVB3- or mock infected mice.

Fig. S4. Reproducibility analysis for DBS sampling and proteomics analysis.

Fig. S5. Protein detectability analysis in proteomics assays used to analyze DBS samples from CVB3 infected and mock infected animals.

Fig. S6. An overview of the interactive web-based interface for proteomics data generated through the analysis of longitudinally collected DBS samples.

Fig. S7. Pancreatic beta cell loss and exocrine pancreas damage in CVB3 infected SOCS-1-tg mice is prevented by an early intervention.

