## Supplementary material for "Longitudinal blood microsampling and proteome monitoring facilitate timely intervention in experimental type 1 diabetes": Figures S1-S7

Anirudra Parajuli *et al.*

**This file includes:** Figs. S1 to S7

**Other Supplementary Materials:** Will be provided upon publication

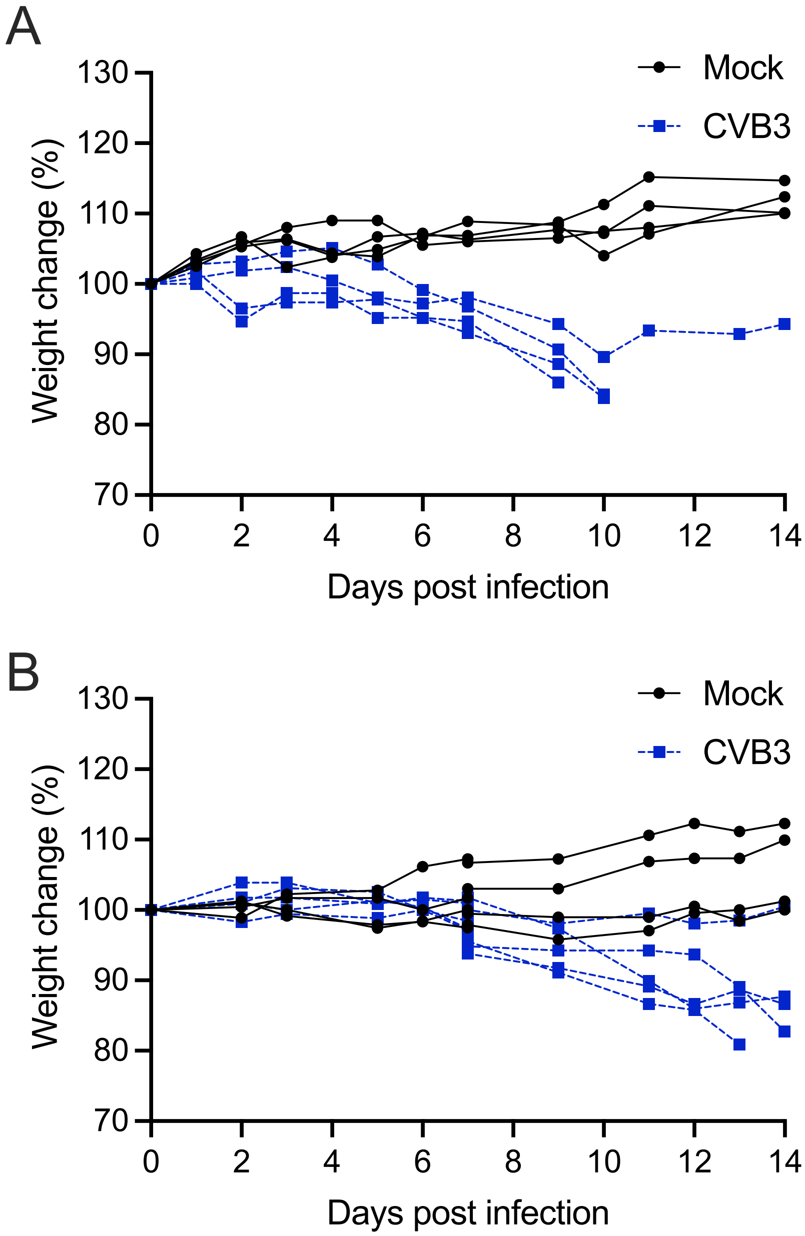


**Fig. S1. Percentage weight change in mock and CVB3-infected NOD mice.** (**A** and **B**) NOD mice aged 8-9 weeks were infected with CVB3 (200 µl RPMI medium containing $10^5$ PFU CVB3, i.p.) or mock infected (200 µl RPMI medium, i.p.). Weight measurements were taken as indicated, and animals were euthanized on day 14 post-infection or when weight loss exceeded 15 % of their maximal weights. The results from two separate studies are shown (A, Study 1 and B, Study 2). Each line represents an individual animal (black lines: mock-infected animals; hatched blue lines: CVB3-infected animals). In panel A: mock-infected animals, n = 4; CVB3-infected animals, n = 4. In panel B: mock-infected animals, n = 4; CVB3-infected animals, n = 5.

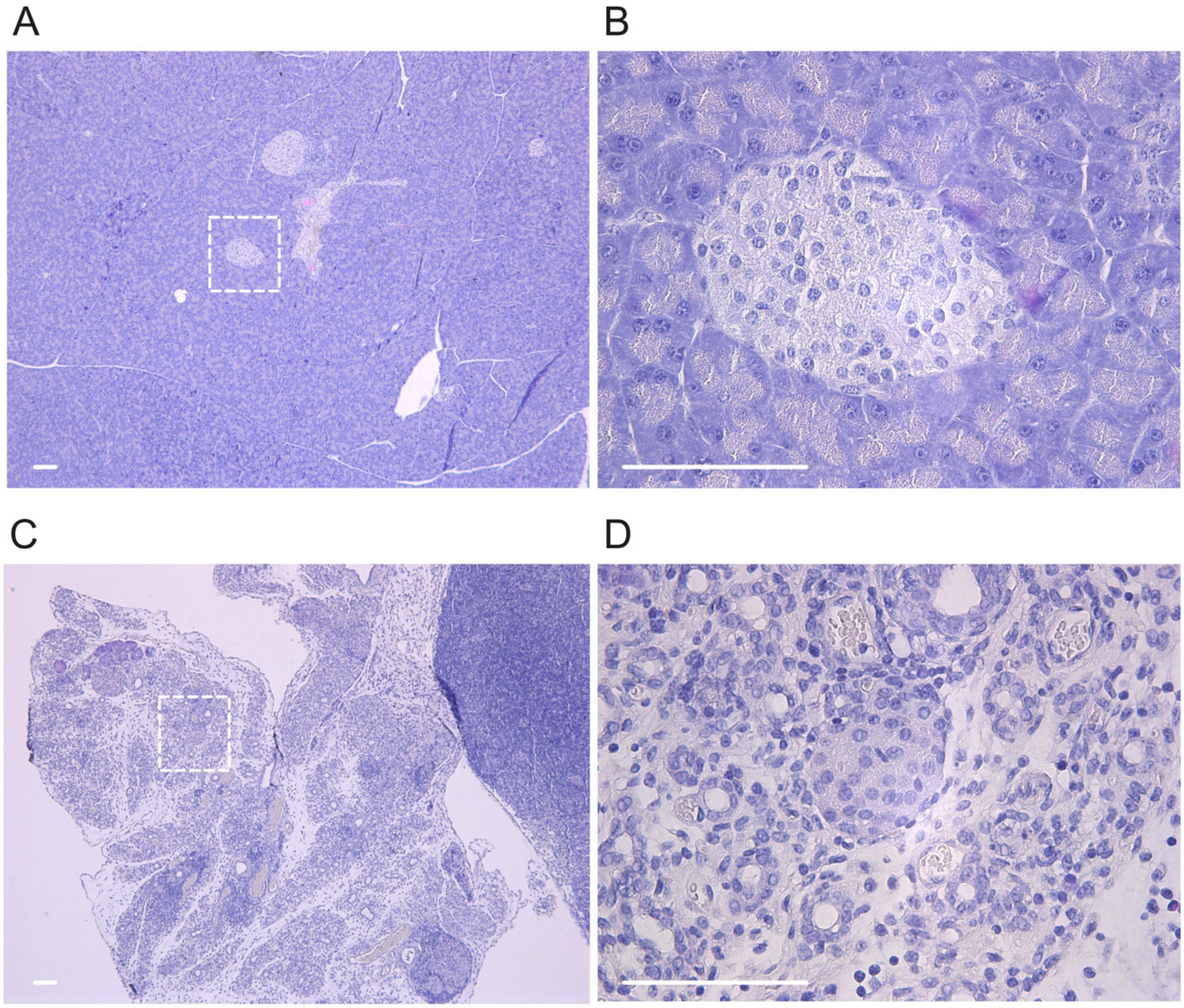


**Fig. S2. NOD mouse pancreas displays clear signs of pathology after CVB3 infection.** NOD mice aged 8-9 weeks were infected with CVB3 (200 µl RPMI medium containing $10^5$ PFU CVB3, i.p.) or mock infected (200 µl of RPMI medium, i.p.). Animals were sacrificed on day 14 p.i. or if their weight loss exceeded >15% of their highest body weight. (**A** and **B**) Representative images from a H&E stained formalin fixed pancreas specimen from a mouse that was mock-infected and sacrificed on day 14 p.i. (**C** and **D**) Representative images of a H&E stained formalin fixed pancreas specimen from a mouse that was infected with CVB3 and sacrificed on day 14 p.i. Images taken at 5x magnification (A and C) and at 40x magnification (B and D). Scale bars = 100 µm.

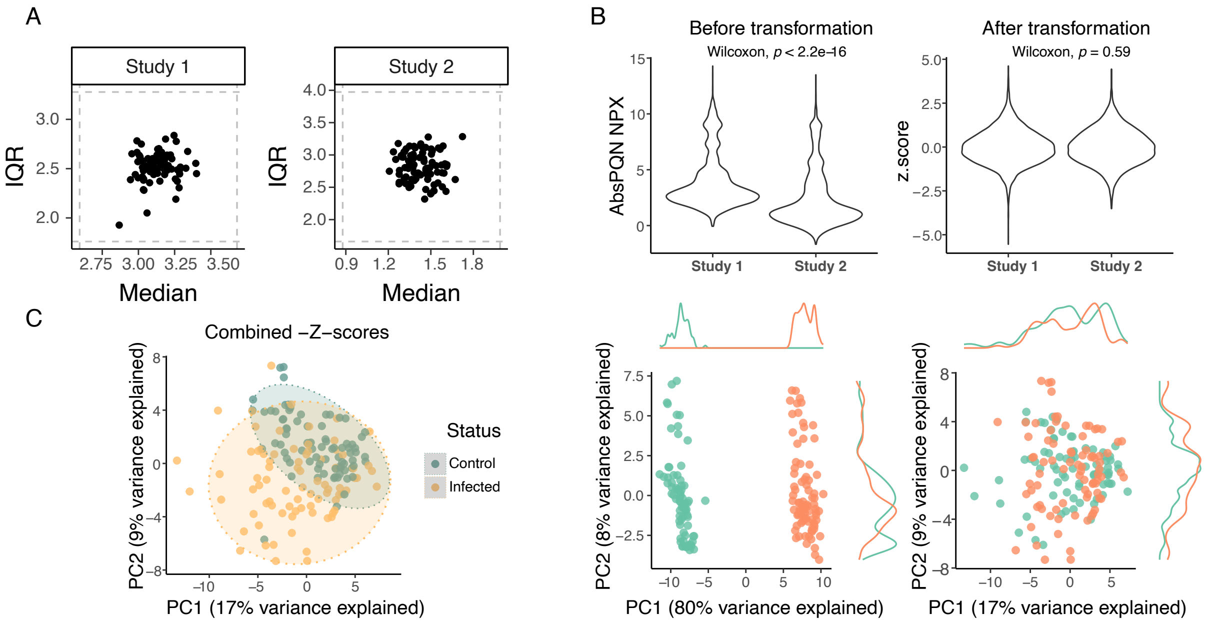


**Fig. S3. Initial data analysis of proteomics data generated from dried blood spot samples collected from CVB3- or mock infected mice.** Dried blood spot samples were collected from CVB3- or mock-infected NOD mice over 14 days post infection in two separate studies (Study 1 and Study 2) as outlined in Fig. 1A. Proteins were measured in the samples using proximity extension assays. (**A**) Outlier analysis: Sample median against sample interquartile range (IQR) for each study. Each point is a sample, and the dotted line represents a threshold of 5 standard deviations from the mean IQR or sample median. Samples outside the threshold may be outlier samples. Here, no samples are outside of the threshold. (**B**) Data processing: Effect of Z-score transformation on the Olink data from the two studies. Overall protein signals (upper panel) for the two studies were significantly different before transformation using Wilcoxon Rank sum test, but not after. The lower panel shows overlapping datasets after transformation. (**C**) Data overview: PCA plot showing the separation between infected and control group for Z-transformed data.

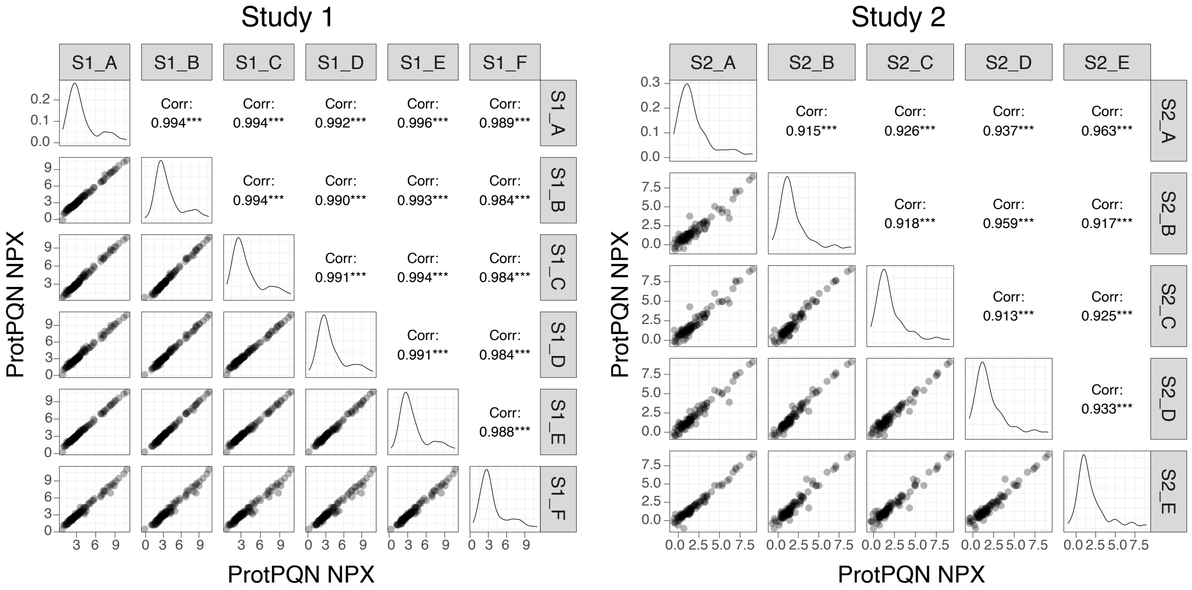


**Fig. S4. Reproducibility analysis for DBS sampling and proteomics analysis.** In two separate studies (Study 1 and Study 2), DBS samples were collected from CVB3- and mock-infected NOD mice over 14 days as indicated in Fig. 1A. At the end of each study, multiple DBS samples were collected from a single mock-infected animal. Proteins in the DBS replicate samples from both studies were measured via proximity extension assays. Shown are Spearman correlations for the DBS samples taken within each study (right panel, Study 1, n = 6; left, Study 2, n = 5).

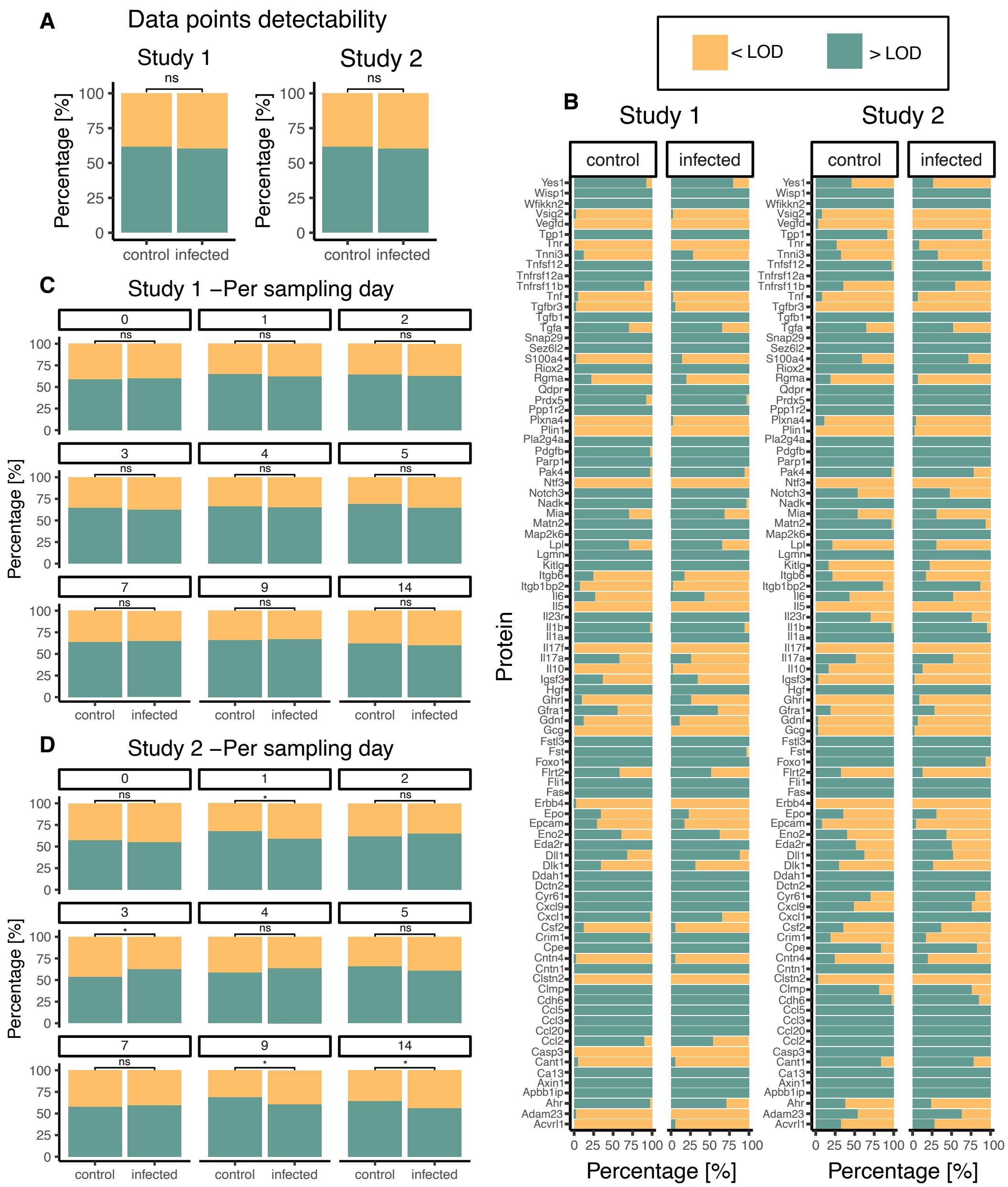


**Fig. S5. Protein detectability analysis in proteomics assays used to analyze DBS samples from CVB3 infected and mock infected animals.** DBS samples were collected from CVB3- or mock-infected NOD mice over 14 days after infection in two separate studies (Study 1 and Study 2) as indicated in Fig. 1 A and proteins were measured in these samples via proximity extension assays (a total of unique 92 proteins were included). The obtained protein signal for each sample was compared to the limit of detection (LOD) for each protein. (**A**) Percentage of data points above and below LOD in infected and control group for Study 1 (left) and Study 2 (right). Green indicates the data points above LOD, and yellow the datapoints below LOD. The differences between the groups were not significant (ns) in either of the studies. (**B**) Detectability per protein and per group for the two studies. (**C** and **D**) Detectability per group and per sampling day for Study 1 and 2, respectively. Significance between groups were calculated using Fisher's Exact test. * $P < 0.05$.

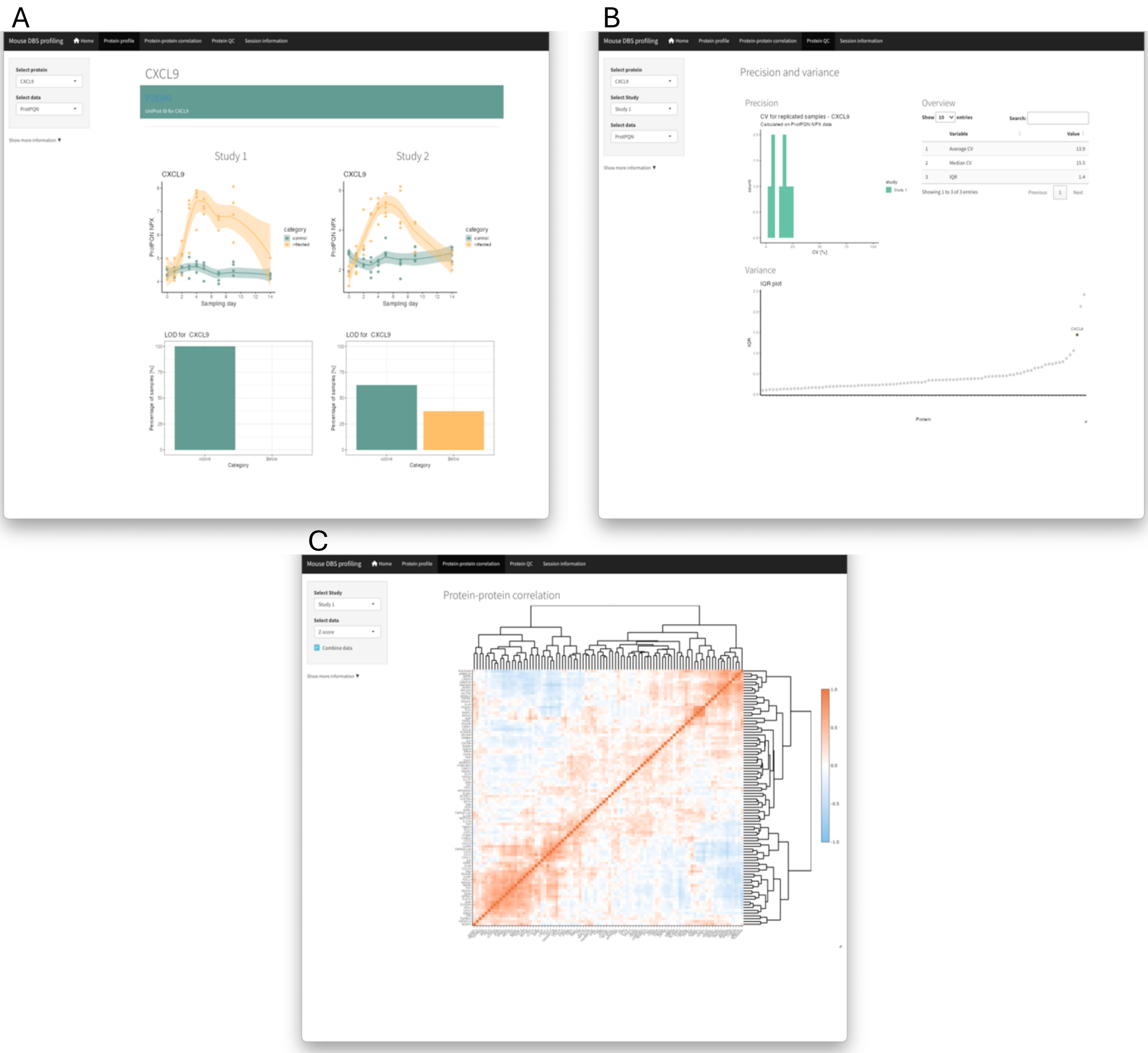


**Fig. S6. An overview of an interactive web-based interface for proteomics data generated through the analysis of longitudinally collected DBS samples.** An interactive web-based interface was developed that contains the proteomics data measured in dried blood spot samples from CVB3- and mock-infected mice. (**A**) The app has a tab showing the protein profiles and detectability for each protein included in the study. (**B**) A second tab shows the CVs and inter-quartile-range for each protein. (**C**) A third tab shows the protein-protein correlations for the combined data or separately for the different study sets. URL will be provided upon publication.

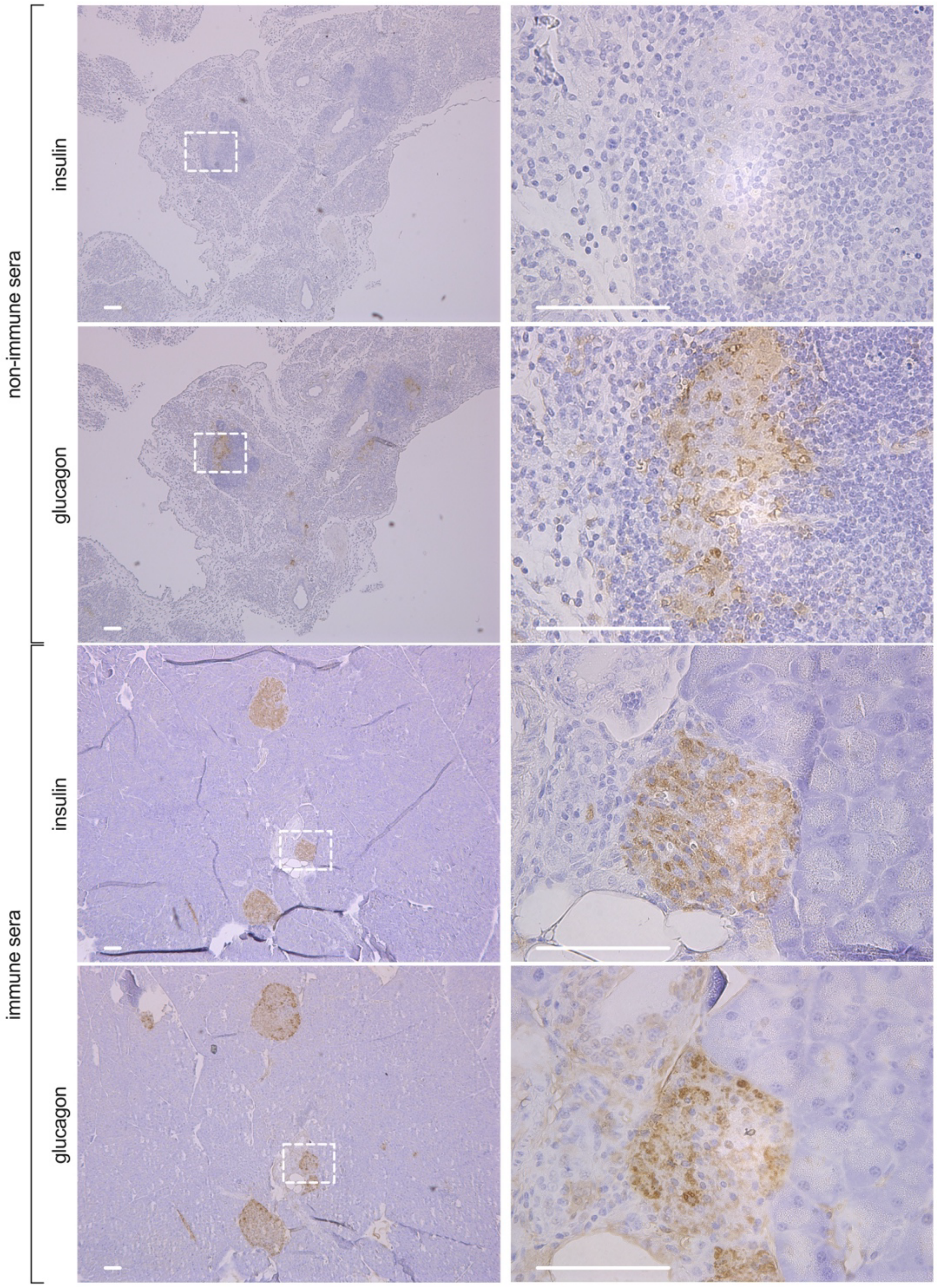


**Fig. S7. Pancreatic beta cell loss and exocrine pancreas damage in CVB3 infected SOCS-1-tg mice is prevented by an early intervention.** SOCS-1-tg mice were infected with CVB3 ($10^5$ PFU CVB3, i.p., n = 14). On days 2 and 3 post infection, animals were treated with either non-immune (n = 8) or immune sera (n = 6). Animals were sacrificed on day 14 p.i. or at diabetes diagnosis. Representative images of insulin or glucagon antibody stained sequential pancreas sections from mice infected with CVB3 and treated with non-immune sera (upper panels) or immune sera (lower panels). Left column shows the wider landscape of the pancreas at 5x magnification, and right column shows the highlighted squares at 40x magnification. Positive areas are stained brown. Scale bar = 100 µm.
